# A live-cell ergosterol reporter for visualization of the effects of fluconazole on a human fungal pathogen

**DOI:** 10.1101/2023.09.14.557744

**Authors:** Antonio Serrano, Miguel A. Basante-Bedoya, Martine Bassilana, Robert A. Arkowitz

## Abstract

Ergosterol, an essential plasma membrane amphipathic lipid, is a major component of the fungal plasma membrane. Most fungal pathogens are sensitive to azole drugs that target ergosterol biosynthesis and resistance/tolerance to azoles is increasingly problematic. <i>Candida albicans</i> is the most prevalent etiology of candidiasis and, in this fungal pathogen, ergosterol rich sub-domains are likely to include sphingolipids, as well as specific membrane proteins, such as multidrug transporters. To investigate the dynamics of ergosterol during the cell cycle and whether drug treatment affects these dynamics in this opportunistic pathogen, we adapted the D4H (domain 4 of the perfringolysin O bacterial toxin) reporter for studying sterol-rich membrane domains. We show that D4H provides a direct readout for the cellular effects of fluconazole and that highly polarized ergosterol is not critical for budding or filamentous growth.

---

Most fungal pathogens are sensitive to azole drugs that target ergosterol biosynthesis. Resistance and tolerance to azoles are particularly problematic because these drugs are used extensively in medical, as well as agricultural, applications. The emergence of new drug resistant fungal variants and species, including *Candida auris*, is a global concern recently recognized by the Word Health Organization and the US Centers for Disease Control (1, 2). The majority of sterols are found in the plasma membrane, with ergosterol making up > 10 mol % of the *Saccharomyces cerevisiae* lipidome (3) and approximately 40 mol % of the plasma membrane lipids (4). In *S. cerevisiae*, ergosterol is found predominantly in the inner leaflet of the plasma membrane and roughly half of all lipids in this leaflet are sterols, approaching their solubility limit (5).

In fungi, ergosterol, an essential plasma membrane amphipathic lipid, is critical for membrane fluidity. *Candida albicans* is the most prevalent etiology of candidiasis and, in this fungal pathogen, ergosterol rich sub-domains are likely to include sphingolipids, as well as specific membrane proteins such as those involved in lipid metabolism and multidrug transporters (6). Interestingly, *C. albicans* mutants with reduced ergosterol levels trigger weaker host immune responses (7), and are defective in the lysis of host macrophages (8, 9). This suggests that sterols are important for interactions with host cells.

An important question is, what are the dynamics of ergosterol during the cell cycle and whether drug treatment affects these dynamics? To address this question, one must be able to visualize the distribution of ergosterol within the plasma membrane, which is challenging. Filipin, a fluorescent polyene macrolide antibiotic, has potent antifungal activity, and induces membrane deformations and lesions (10, 11), limiting its use for live-cell imaging (12, 13). Recently, an improved version of the domain 4 of the perfringolysin O bacterial toxin (14) that binds free membrane sterols down to a threshold of 20 mol %, D4H, was used to visualize ergosterol dynamics in the live cells of model yeasts *Schizosaccharomyces pombe* and *S. cerevisiae* (8, 15, 16). In this study, we adapted this D4H reporter assay for studying sterol-rich membrane dynamics in *C. albicans*, a human fungal commensal and opportunistic pathogen. We show that D4H provides a direct readout for the cellular effects of fluconazole in *C. albicans* and that highly polarized ergosterol is not critical for budding or filamentous growth in this organism.

## Results & Discussion

### Ergosterol dynamics during *C. albicans* growth

D4H is a 110 amino acid polypeptide that specifically binds membrane ergosterol and cholesterol. DNA encoding the D4H domain was fused to that of a codon optimized red fluorescent protein (RFP) (17). The resulting RFP-D4H biosensor was used to probe ergosterol dynamics during budding and hyphal growth. In stationary cells, as well as in cells with sub-optimal growth, punctae of RFP-D4H were observed, typically associated with the cell cortex (Fig. 1A). These punctae partially co-localized with sites of endocytosis, as visualized by the actin-binding protein 1 (Abp1) in double labeling experiments (Fig. S1). In *S. pombe*, sterols also accumulate in endosomal compartments (16), and in *S. cerevisiae* esterified ergosterol is found in lipid droplets (15). Hence we also examined whether these RFP-D4H punctae co-localized with endosomal marker Snf7 or the lipid droplet dye BODIPY, and did not observe co-localization with either of these compartment markers. When active growth on agar pads was maintained during image acquisition, ergosterol was enriched at sites of polarized growth. Upon bud emergence, ergosterol was substantially enriched at the bud tip, concomitant with a decrease in the signal associated with punctae (Fig. 1A), and upon germ tube emergence, a tight cluster of ergosterol was observed at the apex (Fig. 1B) (17). This polarized distribution of ergosterol to sites of growth was maintained throughout the cell cycle, also appearing at cell division sites. Specifically, during either bud or germ tube emergence, ergosterol accumulated at the nascent growth site, prior to a visible protrusion. The tight cluster of ergosterol was localized to the growing tip persistently as the filament extended (17). This polarization of ergosterol at growth sites was observed in the majority (86% and 72%, respectively) of budding and filamentous cells (Fig. 1C).

**Figure 1.**
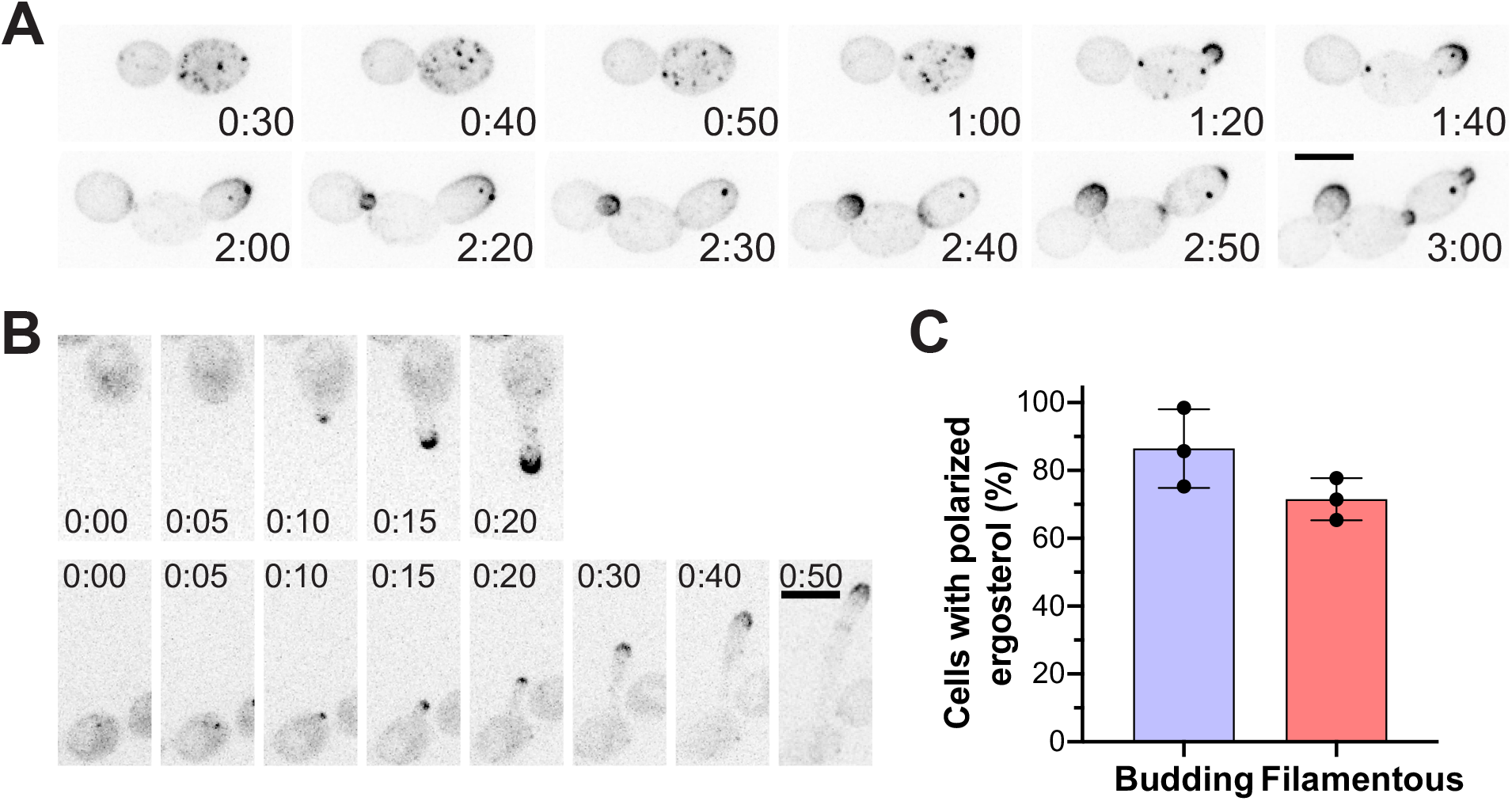
Dynamics of ergosterol during budding and filamentous growth. **A)** Ergosterol distribution is dynamic during the cell cycle. Representative time-lapse images of wild-type cells expressing RFP-D4H (PY6037) at indicated times are shown; images are maximum projections of 26 0.4 µm z-sections. **B)** Ergosterol polarization persists at the filament apex. Representative images from 2 time-lapse experiments of wild-type cells expressing RFP-D4H (PY6037) at indicated times, grown in the presence of FBS at 37ºC, are shown; images are maximum projections of 21 0.4 µm z-sections. **C)** Ergosterol is highly polarized at the growth sites of both budding and filamentous cells. Quantitation of cells with polarized RFP-D4H signal in small buds or in filament tips (3 independent experiments, with *n* = 40-80 cells per experiment).

In *Candida glabrata* cholesterol uptake can occur from the host (18). Hence, we examined whether ergosterol tip enrichment was observed when filamentous growth was induced in Spider media, which does not contain cholesterol. In *C. albicans*, we detected ergosterol apical enrichment in hyphae induced in Spider media; the levels of the D4H reporter at the apex were indistinguishable from that of hyphae induced with serum (Fig. S2). Together, these data indicate that highly polarized ergosterol is present in actively growing cells irrespective of cell cycle stage and growth mode.

### Polarized ergosterol distribution is not critical for growth

Azole drugs inhibit lanosterol demethylase, encoded by the *ERG11* gene, and lead to a reduction in ergosterol. Hence, we next examined the distribution of ergosterol upon inhibition or repression of Erg11. Upon incubation with a high concentration (10X MIC or above) of fluconazole (FCZ) for 16 hr, essentially no cells with polarized D4H signal were observed (Fig. 2A). We verified that the RFP-D4H reporter did not substantially alter fluconazole sensitivity by growth on media containing 10 µg/ml of this drug (Fig. S3A). Interestingly both in budding cells (Fig. 2B) and in filamenting cells (Fig. 2C, 2D) exposed to fluconazole for 2-3 hr, ergosterol was no longer enriched at growth sites. Nonetheless, budding growth continued in the presence of these high fluconazole concentrations, with doubling times indistinguishable from that of control cells (Fig. S3B). Strikingly, within this time frame, filament extension rates were not drastically altered (Fig. S3C). Reduced filament extension rates (by ~40%) were only observed at later times (4 hr). These results are consistent with inhibition of Erg11 dramatically reducing polarized ergosterol, on the order of hours.

**Figure 2.**
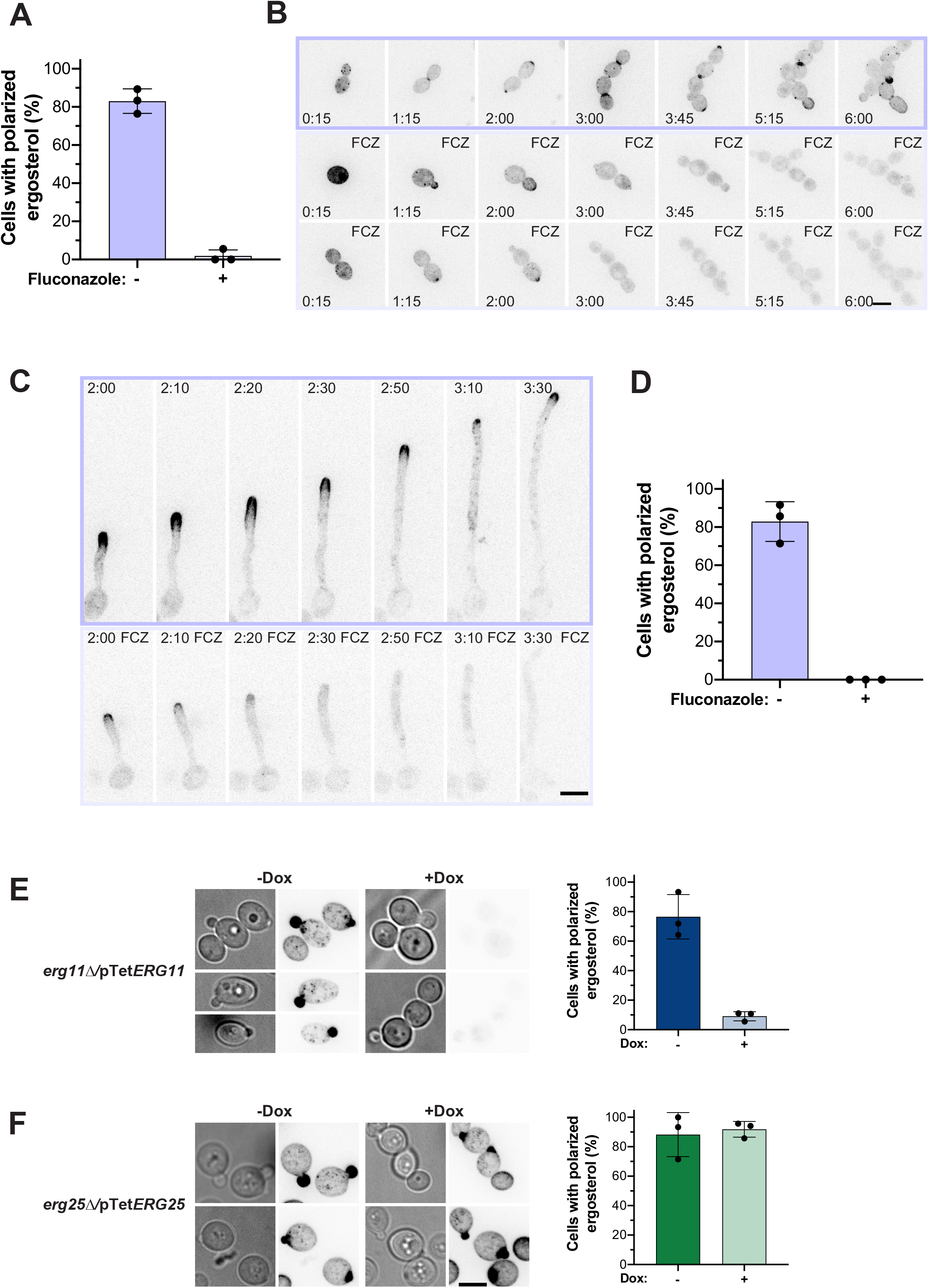
Fluconazole or repression of *ERG11* results in rapid ergosterol depletion at growth sites. **A)** Fluconazole alters ergosterol polarization in budding cells. Wild-type cells expressing RFP-D4H (PY6037) were grown with or without 10 µg/ml fluconazole (FCZ) for 16 hr; the percentage of cells with RFP-D4H signal associated with small buds is shown (3 independent experiments with *n* = 10-30 cells per experiment). Error bars indicate standard deviations. **B)** Ergosterol polarization is reduced upon fluconazole exposure in budding cells. Representative images from time-lapse experiments of wild-type cells expressing RFP-D4H (PY6037), grown with (2 time-lapse experiments) or without FCZ, are shown. **C)** Ergosterol polarization is reduced upon fluconazole exposure in hyphal cells. Representative images from time-lapse experiments of wild-type cells expressing RFP-D4H (PY6037), grown in the presence of FBS with or without FCZ, are shown. **D)** Quantitation of hyphal cells with tip enriched ergosterol. Wild-type cells expressing RFP-D4H (PY6037) were grown with or without FCZ; the percentage of cells with RFP-D4H signal at filament apex at 2 ½ hr incubation time is shown (3 independent experiments, with *n* = 10-14 cells per experiment). Error bars indicate standard deviations. **E)** Polarized distribution of ergosterol depends on Erg11. Representative images of *erg11∆*/pTet*ERG11* cells (PY6862), grown in the absence (-Dox) or presence of doxycycline (+Dox), are shown (left panel). Quantitation of cells with a polarized RFP-D4H signal in small buds is shown (3 independent experiments, *n* = 10-30 cells per experiment) (right panel). Error bars indicate standard deviations. **F)** Polarized distribution of ergosterol does not require Erg25. Representative images of *erg25∆*/pTet*ERG25* cells (PY6859), grown as in Fig. 2E, are shown (left panel). Quantification of cells with a polarized RFP-D4H signal in small buds, as in Fig. 2E (right panel). Error bars indicate standard deviations.

We took a complementary approach to deplete Erg11, using a strain where the sole copy of *ERG11* was behind a doxycycline (Dox) repressible promoter (pTet-*ERG11*). This strain had a substantial growth defect when incubated with Dox (20 µg/ml), which was not altered by the presence of RFP-D4H and RT-PCR analyses did not detect that *ERG11* transcripts after overnight growth in Dox (Fig. S4A, S4B). The inhibition of pTet-*ERG11* expression by doxycycline was nearly complete, as few if any cells had polarized ergosterol in the presence of Dox, compared to 80% in its absence (Fig. 2E). As a control for the effect of doxycycline on D4H localization, we examined whether Dox-dependent depletion of *ERG25* altered the polarized distribution of ergosterol using a strain, in which the sole copy of *ERG25* was under the control of the repressible promoter. Adding Dox to this strain resulted in reduced *ERG25* transcript levels (Fig. S4C), yet the percentage of cells with polarized ergosterol during budding growth was similar in the absence and presence of Dox (Fig. 2F). This indicates that the D4H domain does not bind sterol precursors or aberrant sterols generated in bypass pathways, consistent with the idea that this domain specifically binds ergosterol and does not bind lanosterol (19).

In summary, our study of a specific live-cell ergosterol reporter for *C. albicans* reveals that apical enrichment of this sterol is not critical for budding and filamentous growth in this human fungal pathogen. Furthermore, it highlights that this live-cell reporter of ergosterol localization is likely to be a useful tool in the analyses of azole resistance and tolerance mechanisms, including alterations in drug targets and upregulation of efflux activities.

## MATERIAL AND METHODS

### Strains and media

Strains used in this study are listed in Table S1. For transformation, strains were grown in YEPD (yeast extract, peptone, dextrose) supplemented with Uridine (80 µg/ml) at 30ºC. For budding growth experiments, cells were grown in synthetic complete (SC) medium, supplemented with Uridine at 30ºC. For filamentous growth, induction of cells was carried out on agarose pads with either 75% fetal bovine serum (FBS; PAN Biotech) in SC medium or with Spider medium at 37ºC. For doxycycline (Dox) gene repression, SC was supplemented with 20 µg/ml Dox (2). For fluconazole (FCZ) experiments, cells were grown in the presence of 10 µg/ml FCZ, 0.02% DMSO or with 0.02% DMSO as a control. Lipid droplets were stained with BODIPY (ThermoFisher Sci). Cells were grown overnight in SC media and incubated with 10 µM BODIPY for 30 min at room temperature prior to imaging.

The oligonucleotides used in this study are listed in Table S2. To visualize the distribution of ergosterol, we used the genetically encoded biosensor D4H with the plasmid *pExp-pACT1-mScarlet-D4H* (17). This plasmid was linearized with NcoI and integrated into the *RP10* locus. The Dox repressible *erg11Δ/pTetERG11* and *erg25Δ/pTetERG25* strains were constructed from PY173, a derivative of BWP17 containing the tetracycline-regulatable transactivator TetR-ScHAP4AD, as described (20). The Abp1-GFP and Snf7-GFP strains were generated by homologous recombination, using pFA-GFPγ-URA (21) or pGFP-NAT1 (22) and primer pairs ABP1.P1/ABP1.P2 and SNF71.P1/SNF7.P2, respectively.

### Microscopy and image analyses

Cells were imaged as described previously using spinning-disk confocal microscopy (17, 23) with a PLANAPO total internal reflection fluorescence (TIRF) 1.45-numerical-aperture (NA) 100× objective. All live-cell imaging timelapses were performed starting with washed overnight grown cells incubated on agarose pads. To quantify D4H associated signal intensity at growth sites, sum projections of z-stacks were analysed using Volocity software version 6.3 (Perkin Elmer, USA) and signal was considered polarized if it was 5 or 7 standard deviations above the image mean in budding and filamentous cells, respectively. Cell doubling times were determined from time-lapse images, from bud emergence to subsequent bud emergence. D4H associated signal intensities at filament tips were determined from exponentially grown cells incubated for 1-2 hr either in Spider (24) or FBS containing media. The apex/cytoplasm ratio of D4H signal was determined from image central z-sections, quantified with Fiji (Version 1.54f), using regions of interest (ROI) at the tip of the filamentous cells (apex) and 1-2 µm from this area (cytoplasm). Ratios of 1.7 or greater were scored as polarized.

### RNA extraction and RT-PCR

Cells were incubated overnight with SC media in the absence or presence of 20 µg/ml of Dox. mRNA extraction and RT-PCR were carried out as described (17). Oligonucleotide pairs ACT1.P1/ACT1.P2, ERG11.P1/ERG11.P2 and ERG25.P1/ERG25.P2 were used to amplify *ACT1, ERG11* and *ERG25*, respectively.

### Statistical analysis

Data were compared by unpaired *t-test* using GraphPad Prism (v. 8) software, with all *p* values indicated in figure legends.

## Figure legends

**Figure S1. Cortical membrane associated ergosterol partly co-localizes with endocytosis sites. A)** Ergosterol punctae partly co-localize with endocytosis sites. Overnight cultures of a strain (PY7044) expressing RFP-D4H (magenta) and Abp1-GFP (green) were imaged. Central z-sections are shown. **B)** Ergosterol punctae do not co-localize with the endosomal compartment. A strain (PY7014) expressing RFP-D4H (magenta) and Snf7-GFP (green) was imaged as in S1A and maximum projections of z-sections are shown. **C)** Ergosterol punctae do not co-localize with lipid droplets. A strain (PY6037) expressing RFP-D4H (magenta), with Bodipy labeled lipid droplets (green) was imaged as in S1A and maximum projections of z-sections are shown. Bar, 5 µm.

**Figure S2. Ergosterol enrichment at the hyphal apex is independent of the hyphal growth inducer**. Representative images of wild-type cells expressing RFP-D4H (PY6037), grown in either FBS or Spider containing media at 37ºC for 90 and 120 min, respectively (left panel). Quantification of the ergosterol enrichment at the filament apex (ratio of the mean D4H apex signal to the cytoplasm; ~2 µm back the apex), *n* = 30-40 cells. Error bars indicate standard deviations. A two-tailed *t-test* revealed no significant difference (ns).

**Figure S3. The ergosterol reporter does not alter fluconazole sensitivity, and growth is not substantially affected by short exposure to fluconazole. A)** The RFP-D4H reporter does not alter fluconazole sensitivity. Serial dilutions of indicated strains (WT, PY4861 and WT expressing RFP-D4H, PY6037) were spotted on YEPD with or without FCZ (10 µg/ml) and incubated for 3 days at 30ºC. **B)** Initial doubling time of cells is similar in the presence and absence of fluconazole. Wild-type cells expressing RFP-D4H (PY6037) were followed by time-lapse microscopy as in Fig. 2A and doubling time from *n* = 70-90 cells was determined. Error bars indicate standard deviations. A two-tailed *t-test* revealed no significant difference (ns). **C)** Initial extension rate of filamentous cells is similar in the presence and absence of fluconazole. Wild-type cells expressing RFP-D4H (PY6037) were analyzed by time-lapse microscopy as in Fig. 2C and filament length in the initial 6 times points (2:00 to 2:50) was fit with a linear regression (r^2^ > 0.95). Extension rates from *n* = 35 cells (from 3 independent experiments) was determined. Error bars indicate standard deviations. A two-tailed *t-test* revealed no significant difference (ns).

**Figure S4. *ERG* gene expression is reduced upon repression of *ERG11* or *ERG25*. A)** The RFP-D4H reporter did not alter sensitivity of the *erg11*Δ/pTet*ERG11* (PY6862) strain to doxycycline. Serial dilutions of wild-type (PY173), *erg11*Δ/pTet*ERG11* (PY6743), and *erg11*Δ/pTet*ERG11* expressing RFP-D4H (PY6862) strains were spotted on YEPD with or without Dox and incubated for 3 days at 30ºC. **B)** RT-PCR was carried out on the *erg11*Δ/pTet*ERG11* (PY6862) strain grown with or without Dox, using a CaERG11TM1 primer pair. Values are the means of determinations from three independent replicates, normalized to *ACT1* and the average level of *ERG11*/*ACT1* in the absence of Dox was set to 1. Error bars indicate standard deviations. A two-tailed *t-test* revealed a significant difference, ***, *P* < 0.001. **C)** RT-PCR was carried out on the *erg25*Δ/pTet*ERG25* (PY6859) strain as in Fig. S4B, using a CaERG25TM1 primer pair. A two-tailed *t-test* revealed a significant difference, *, *P* < 0.04.

**Movie S1**. Ergosterol dynamics in budding cells. Maximum projections of RFP-D4H signal over time.

**Movie S2**. Ergosterol dynamics in filamentous cells. Maximum projections of RFP-D4H signal over time in cells with FBS.

**Movie S3**. Effect of fluconazole on ergosterol dynamics in budding cells. Maximum projections of RFP-D4H signal over time in cells grown with or without 10 µg/ml FCZ.

**Movie S4**. Effect of fluconazole on ergosterol dynamics in filamentous cells. Maximum projections of RFP-D4H signal over time in cells grown in FBS with or without 10 µg/ml FCZ.

## Acknowledgements

We thank S. Martin, S. Bates, and A. Mitchell for strains and plasmids. We thank S. Bogliolo for assistance. We thank J. Berman for comments on the manuscript. We thank the Platforms Resources in Imaging and Scientific Microscopy facility (PRISM; B. Monterroso and S. Ben-Aicha) and Microscopy Imaging Cytometry Côte d’Azur (MICCA) for microscopy support. This work was supported by the CNRS, INSERM, Université Côte d’Azur, ANR (ANR-19-CE13-0004-01), EC Marie Curie IF (101029870) and EC ERC Synergy (951475) grants.

---

**Table S1.**
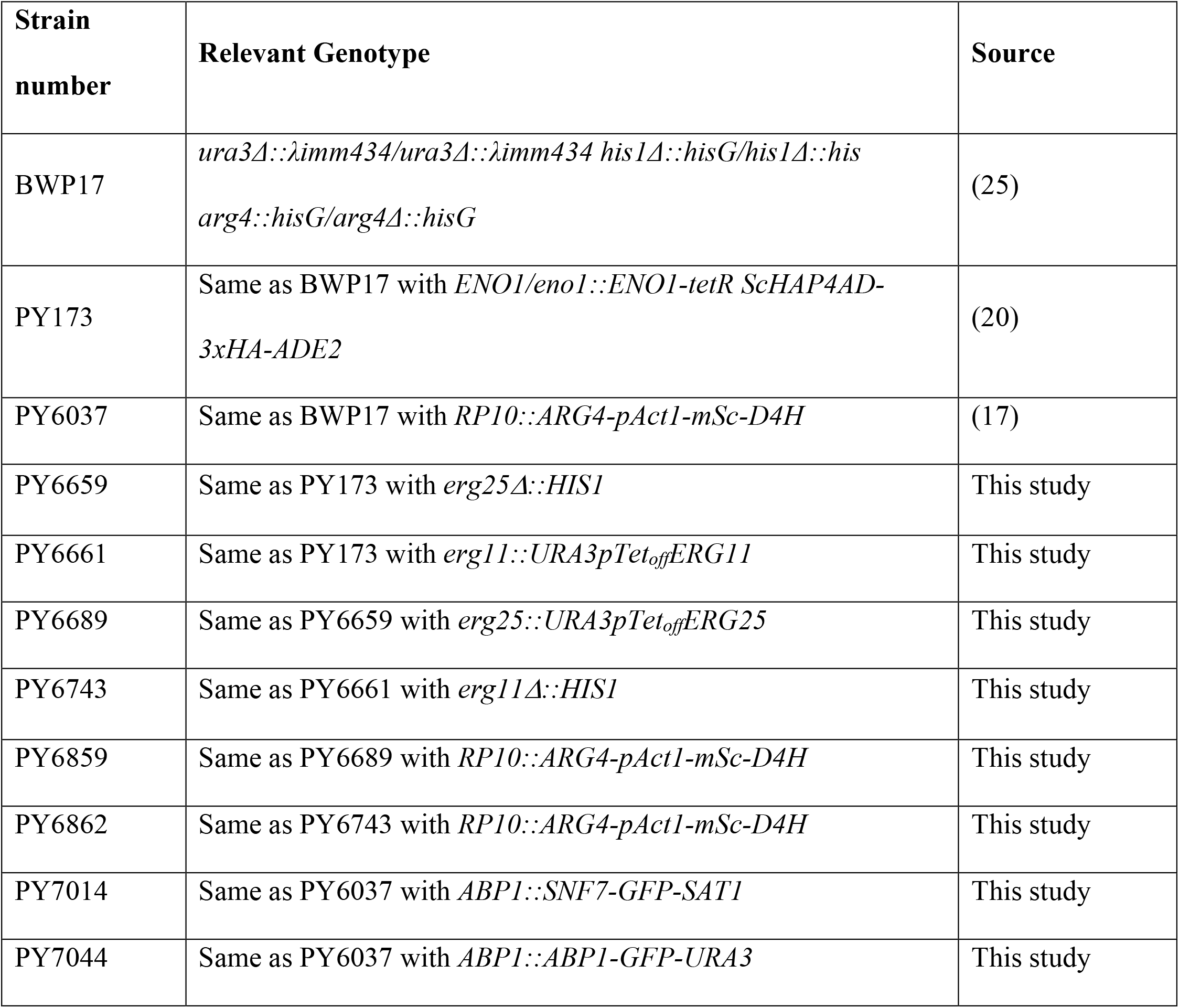
Strains used in this study.

**Table S2.**
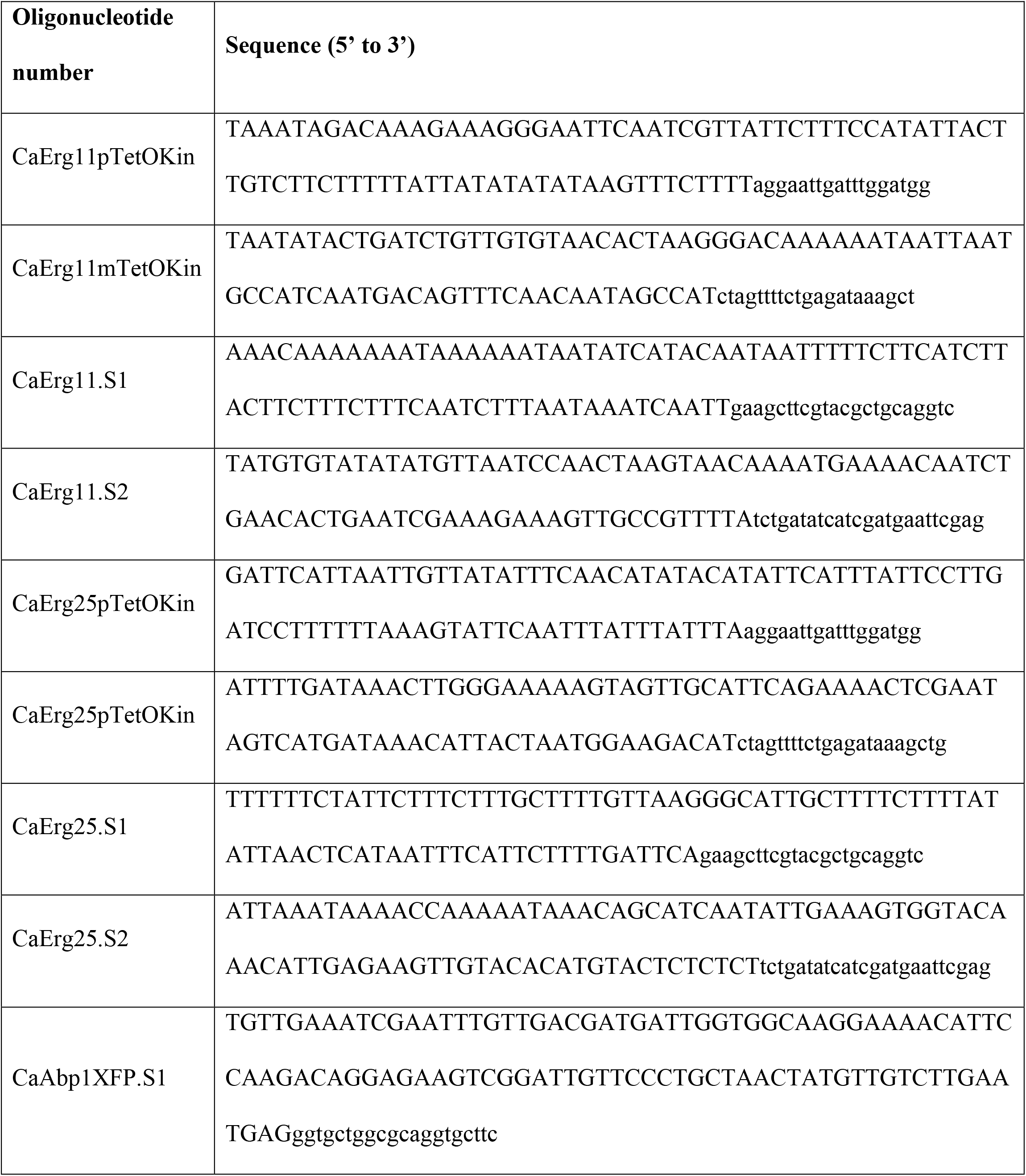
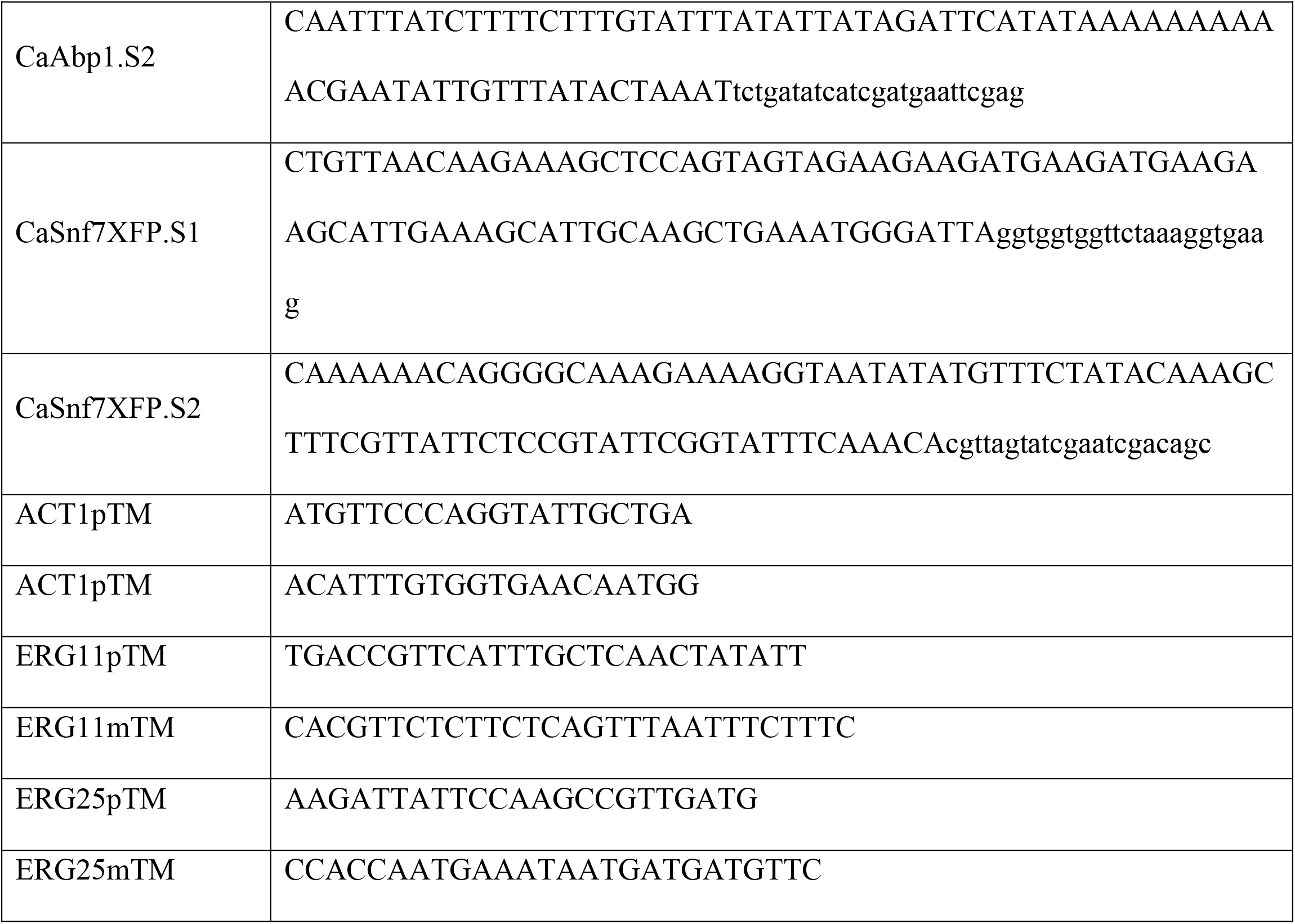
Oligonucleotides used in this study.

