## Supplementary figures and images for "A live-cell ergosterol reporter for visualization of the effects of fluconazole on a human fungal pathogen"

### Fig. S1

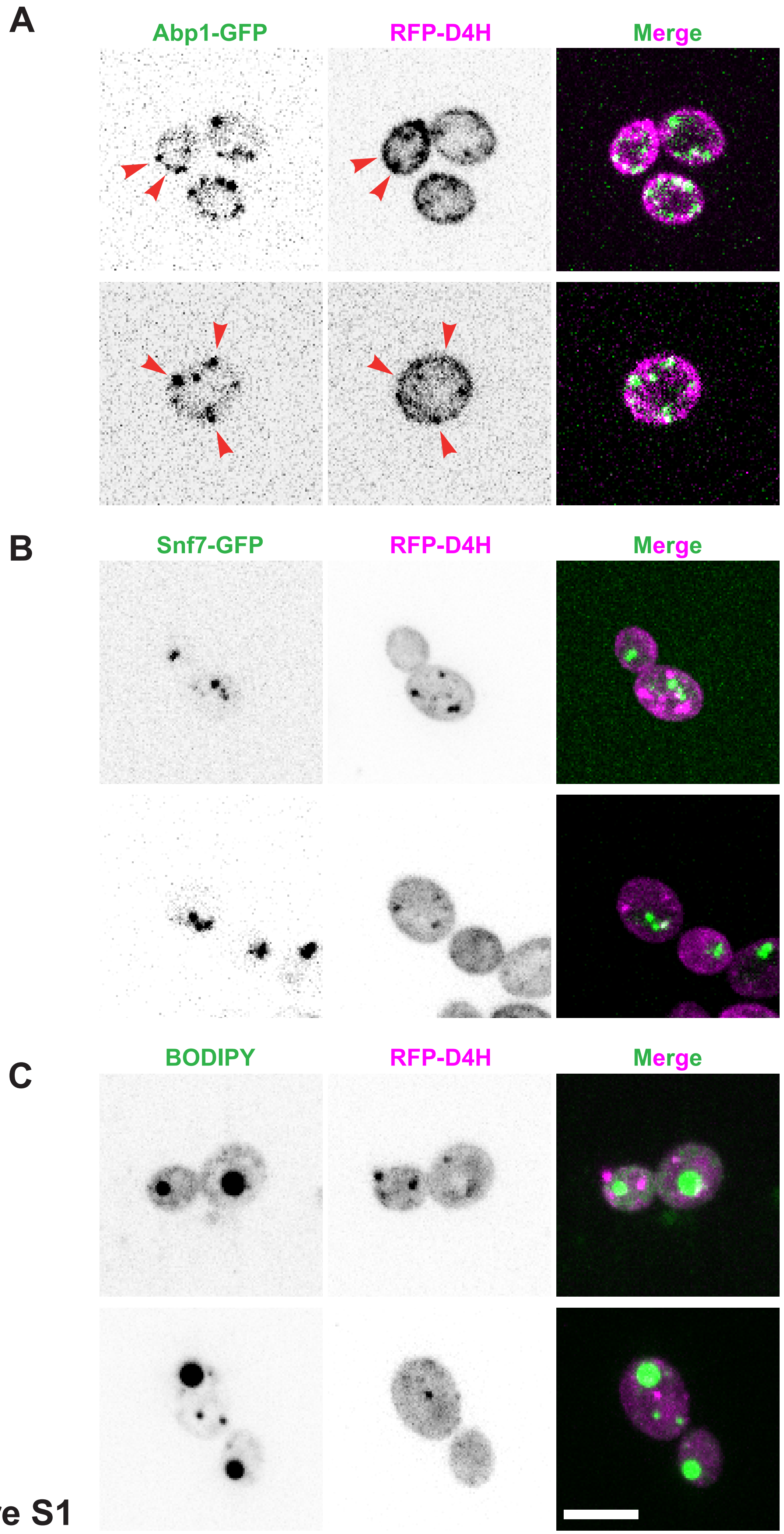

### Fig. S2

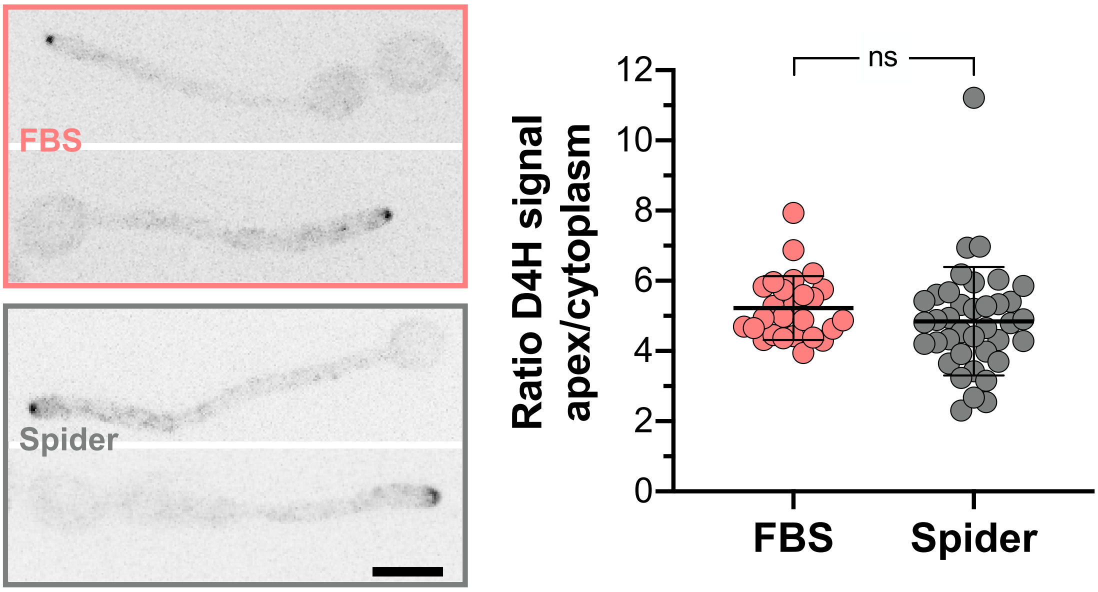

### Fig. S3

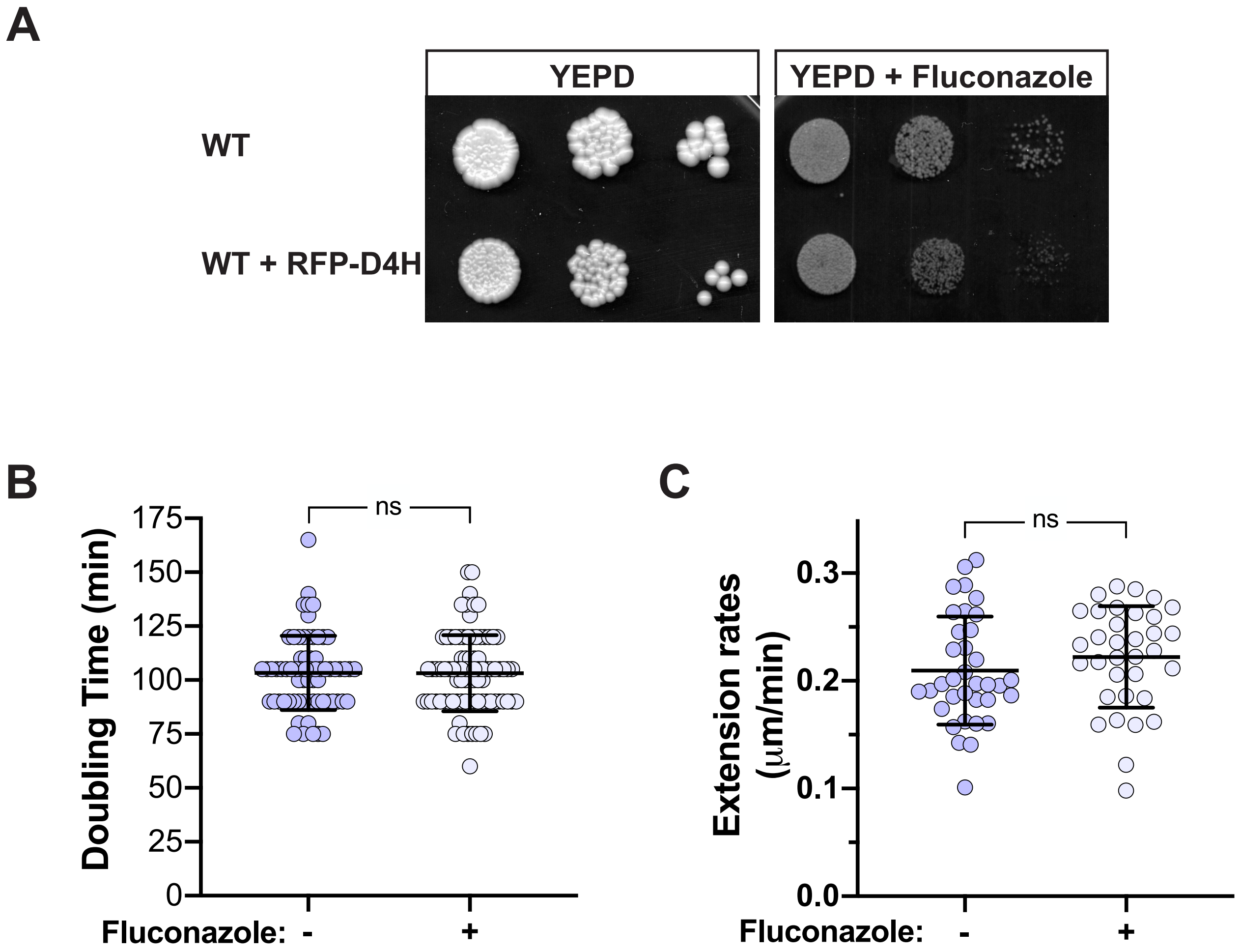

### Fig. S4

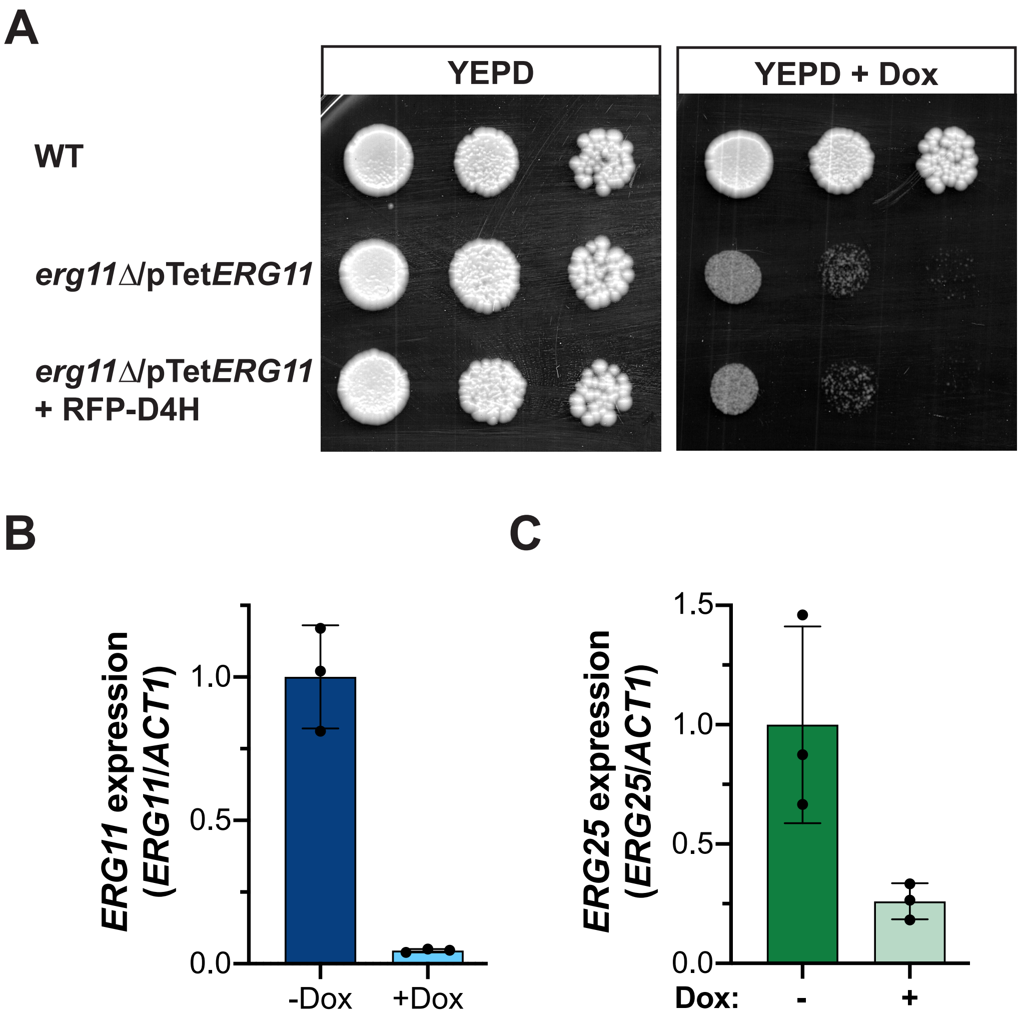
